# Quantitative Models of Lipid Transfer and Membrane Contact Formation

**DOI:** 10.1101/2022.01.17.476625

**Authors:** Yongli Zhang, Jinghua Ge, Xin Bian, Avinash Kumar

**Affiliations:** Department of Cell Biology, Yale University School of Medicine, New Haven, CT, USA; Department of Molecular Biophysics and Biochemistry, Yale University, New Haven, CT, USA; Department of Neuroscience, Yale University School of Medicine, New Haven, CT, USA; State Key Laboratory of Medicinal Chemical Biology, College of Life Sciences, Nankai University, Tianjin, China

**Keywords:** Lipid transfer protein (LTP), membrane contact site (MCS), extended synaptotagmin (E-Syt), optical tweezers, DNA origami, lipid exchange, osmotic membrane tension, VPS13, ATG2

## Abstract

Lipid transfer proteins (LTPs) transport lipids between different organelles in the cell, and thus play key roles in lipid homeostasis and organelle dynamics. The lipid transfer process often occurs at the membrane contact sites (MCS), where two membranes are held within 10-30 nm by LTPs and other membrane tethering factors. While most LTPs act as a shuttle to transfer lipids via their lipid-binding cavities, recent experiments reveal a new category of eukaryotic LTPs that may serve as a bridge to transport lipids in bulk at MCSs, leading to membrane expansion involved in organelle biogenesis. However, the molecular mechanisms underlying lipid transfer and membrane contact formation are not well understood. Here, we first review two recent studies of extended synaptotagmin (E-Syt)-mediated membrane binding and lipid transfer using optical tweezers and DNA origami, respectively. Then we describe mathematical models to quantify the kinetics of lipid transfer by shuttle LTPs based on a lipid exchange mechanism. We find that simple lipid mixing among similar membranes and/or lipid partitioning among different membranes can explain lipid transfer against a concentration gradient widely observed for LTPs. Based on these calculations, we hypothesize that lipid exchange is a general mechanism for lipid transfer by shuttle LTPs. We predict that selective transport of lipids, but not membrane proteins, by bridge LTPs leads to osmotic membrane tension in analog to the osmotic pressure across a semipermeable membrane. A gradient of such osmotic membrane tension and the conventional membrane tension may drive bulk lipid flow through bridge LTPs at a speed consistent with the fast membrane expansion observed *in vivo*. Finally, we discuss the implications of membrane tension and lipid transfer in organelle biogenesis and membrane morphologies. Overall, the quantitative models may help clarify the mechanisms of MCS formation by E-Syts and lipid transfer by LTPs in general.

## Introduction

Lipid transfer proteins (LTPs) specifically transfer lipids between cellular membranes more rapidly than vesicular transport. Consequently, LTPs play key roles in lipid homeostasis, organelle biogenesis, metabolite or ion exchange, and signaling between organelles (Wong *et al*., 2017; Wong *et al*., 2019; Leonzino *et al*., 2021; Reinisch and Prinz, 2021). For example, numerous LTPs mediate cholesterol transport in humans. Dysfunction of LTPs has been associated with various diseases or disorders, including atherosclerosis caused by cholesterol deposition in blood vessels, obesity, and neurodegeneration (Hamilton and Deckelbaum, 2007; Potter *et al*., 2009; Sandhu *et al*., 2018; Ugur *et al*., 2020).

So far, hundreds of LTPs have been discovered, and more are being identified (Wong *et al*., 2019; Hanna *et al*., 2021). Most LTPs contain hydrophobic lipid-binding cavities that specifically bind one or more species of lipids. These LTPs extract lipids from one membrane, hold them in the cavities, diffuse across the cytosol, and insert the lipids into another membrane, thus acting as a shuttle to transfer lipids. A representative shuttle LTP is the *o*xy*s*terol-binding *h*omology protein 4 (Osh4p) in yeast that transfers ergosterol from the endoplasmic reticulum (ER) to the *trans*-Golgi and phosphatidylinositol-4-phosphate (PI4P) backward in a coupled manner (de Saint-Jean *et al*., 2011). Structural, biophysical, and biochemical studies reveal that Osh4p binds ergosterol and PI4P in overlapping cavities and exchanges the two lipids upon binding to membranes (Delfosse *et al*., 2020). Intriguingly, Osh4p can transfer ergosterol against its gradient at the expense of a downgradient transfer of PI4P (de Saint-Jean *et al*., 2011; von Filseck *et al*., 2015b). This counter-exchange activity has been observed both in the cell and in the reconstituted systems. Importantly, the activity is also shared by other Osh proteins and oxysterol-binding protein-related proteins (ORPs) in higher eukaryotes. For example, Osh6, ORP5, and ORP8 counter-transport phosphatidylserine (PS) and PI4P at the ER-plasma membrane (PM) MCSs (Chung *et al*., 2015; Ghai *et al*., 2017; Sohn *et al*., 2018). Yet, the molecular mechanism of LTP-mediated counter-transport is not well understood.

Recently, a new category of eukaryotic LTPs has been discovered that potentially act as bridges to move lipids from ER to other organelles for their biogenesis (Kumar *et al*., 2018; Osawa *et al*., 2019; Valverde *et al*., 2019; Li *et al*., 2020; Melia *et al*., 2020; Hanna *et al*., 2021; Leonzino *et al*., 2021; Reinisch and Prinz, 2021; Wang *et al*., 2021a). These LTPs include ATG2, VPS13, and Ship164, all of which comprise a characteristic chorein motif at their N-termini. Structural studies revealed that these LTPs are club-like and contain a continuous groove 10-20 nm long capped with the chorein domain at one end (Valverde *et al*., 2019; Li *et al*., 2020; Hanna *et al*., 2021). The groove is lined with hydrophobic residues and can bind more than ten lipids in nearly nonspecific manner. Substituting the hydrophobic residues in the middle of the VPS13 bridge with hydrophilic residues impair yeast sporulation (Li *et al*., 2020). In addition, these LTPs contain domains that interact with other proteins residing in the two membranes to help engage the two ends of bridges to membranes (Leonzino *et al*., 2021). All these observations imply that ATG2 and VPS13 may act as a bridge to transport lipids in bulk between membranes. However, a direct test of bulk lipid flow remains missing. A key open question is what drives such directional lipid flow (Reinisch and Prinz, 2021). It has been hypothesized that certain ATPases may pump lipids into the bridges in a manner similar to the lipopolysaccharide-transport protein complex in bacteria (Owens *et al*., 2019). While this hypothesis is appealing, any candidate for such ATPases remains elusive. Therefore, other potential driving forces need to be explored.

Most LTPs transfer lipids at the MCSs (Saheki and De Camilli, 2017a; Wong *et al*., 2019; Prinz *et al*., 2020; Reinisch and Prinz, 2021). Many LTPs contain membrane-binding domains, such as C2, PH, and PX domains, that help tether two membranes to form MCSs (Lemmon, 2008). These domains bind membranes by recognizing specific lipids, Ca^2+^, or other ligands. Consequently, MCSs are highly dynamic and triggered by many signaling clues in the cell, particularly PI(4,5)P2 (Prinz *et al*., 2020). Eukaryotic extended-synaptotagmins (E-Syts), with three isoforms E-Syt1, E-Syt2, and E-Syt3, are representative LTPs and membrane tethers (Min *et al*., 2007; Chang *et al*., 2013; Giordano *et al*., 2013; Saheki and De Camilli, 2017b). They contain an N-terminal hydrophobic hairpin anchored to the ER membrane and a synaptotagmin-like mitochondrial lipid-binding protein (SMP) domain as the lipid transfer module (Saheki *et al*., 2016; Yu *et al*., 2016) (Figure 1A). Two SMP domains form a stable dimer that mediates homo- or hetero-dimerization between E-Syts. The crystal structure of an E-Syt2 fragment reveals two lipids bound by each SMP domain (Schauder *et al*., 2014). A defining feature of E-Syts is an array of C2 domains at the C-termini, five in E-Syt1 and three in E-Syt2 and E-Syt3, connected by disordered polypeptides (Min *et al*., 2007; Tunyasuvunakool *et al*., 2021). Cellular imaging shows that, at a low cytosolic Ca^2+^ level, E-Syt2 and E-Syt3 are constitutively localized at the ER-PM MCSs, whereas E-Syt1 appears diffusive in the entire ER membrane (Giordano *et al*., 2013). An elevation of the cytosolic Ca^2+^ level triggers the recruitment of E-Syt1 to the MCSs, a decrease in membrane separation (Fernandez-Busnadiego *et al*., 2015), and accompanying lipid transfer mediated by all E-Syts.

**Figure 1.**
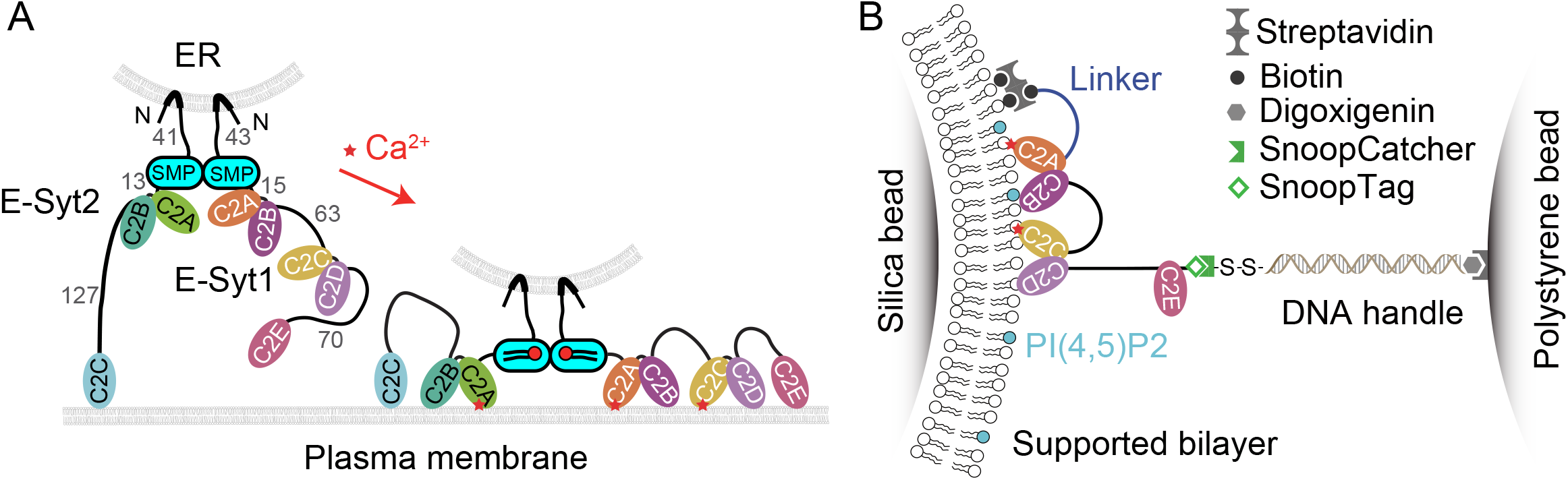
Schematics of E-Syt structures and experimental setup to study C2-membrane binding. A: Domain organization of E-Syts and their roles in forming membrane contact sites (MCSs) and transferring lipids. The Synaptotagmin-like mitochondrial-lipid-binding (SMP) domain mediates dimerization of E-Syts and lipid transfer. The numbers near the disordered polypeptide linkers indicate their length in amino acids. B: Experimental setup to pull the E-Syt1 C2 array (E-Syt1 C2ABCDE) using optical tweezers. The array is attached to the lipid bilayer supported on a silica bead at its N-terminus and conjugated at its C-terminus to a 2,260 bp DNA handle that is immobilized on a polystyrene bead at another end. The extension of the protein-DNA tether is measured to follow dynamic C2-membrane binding and unbinding in the presence of controlled force.

In the following, we first review recent progress in understanding the membrane binding of E-Syts, its role in membrane contact formation, and the mechanism of lipid transfer by the SMP domain. Then, we propose quantitative models of lipid transfer by shuttle LTPs based on the lipid exchange mechanism and test the model with experimental data. Finally, we discuss the roles of canonical and osmotic membrane tension in bulk lipid transfer mediated by bridge LTPs.

### Single-molecule measurements of E-Syt-membrane binding and modeling of MCSs

While the membrane-binding properties of individual C2 domains have been extensively studied (Corbalan-Garcia and Gomez-Fernandez, 2014), it was challenging to characterize the membrane binding of E-Syts containing an array of protein domains that may bind membranes and each other (Bian *et al*., 2018). Furthermore, the membrane binding *in trans* generates a pulling force to the C2 domain, which in turn modulates its membrane binding (Ma *et al*., 2017). To characterize the force-dependent interactions, we applied optical tweezers to dissect the membrane binding of E-Syt1 and E-Syt2 as a function of Ca^2+^ and lipid compositions (Ge *et al*., 2021). A fragment of the C2 containing region of E-Syt1, e.g., E-Syt1 C2ABCDE, is tethered to a lipid bilayer coated on a silica bead at its amino terminus and to a polystyrene bead at its carboxy terminus via a 2,260 bp DNA handle (Figure 1B). The two micron-sized beads are optically trapped with high-resolution dual-trap optical tweezers (Zhang *et al*., 2013) and pulled to apply force to the C2 repeat. The extension of the protein-DNA tether is detected to report dynamic binding and unbinding of the C2 domains to the membrane or other C2 domains (Figure 2A). The E-Syt1 protein undergoes fast and three-state transitions at a constant mean force in the range of 3.6-4.6 pN (Figure 2B), indicative of sequential membrane binding and unbinding of the C2E and C2CD domains (Figure 2A). The membrane binding of the C2AB domain is weak and only observed in the presence of the higher 20 mol% DOPS. The C2CD domain binds to the membrane in a Ca^2+^-dependent manner, while the C2E domain shows moderate membrane affinity in a PI(4,5)P2-dependent but Ca^2+^-independent manner. Pulling a single monomer within the cytoplasmic domain of the E-Syt1 dimer reveals similar membrane binding kinetics, suggesting minimum interactions between the SMP dimer and the membrane. The single-molecule assay did not reveal any significant interactions between different E-Syt1 domains, indicating that these interactions are weak. Similarly, the membrane binding kinetics of E-Syt2 is measured.

**Figure 2.**
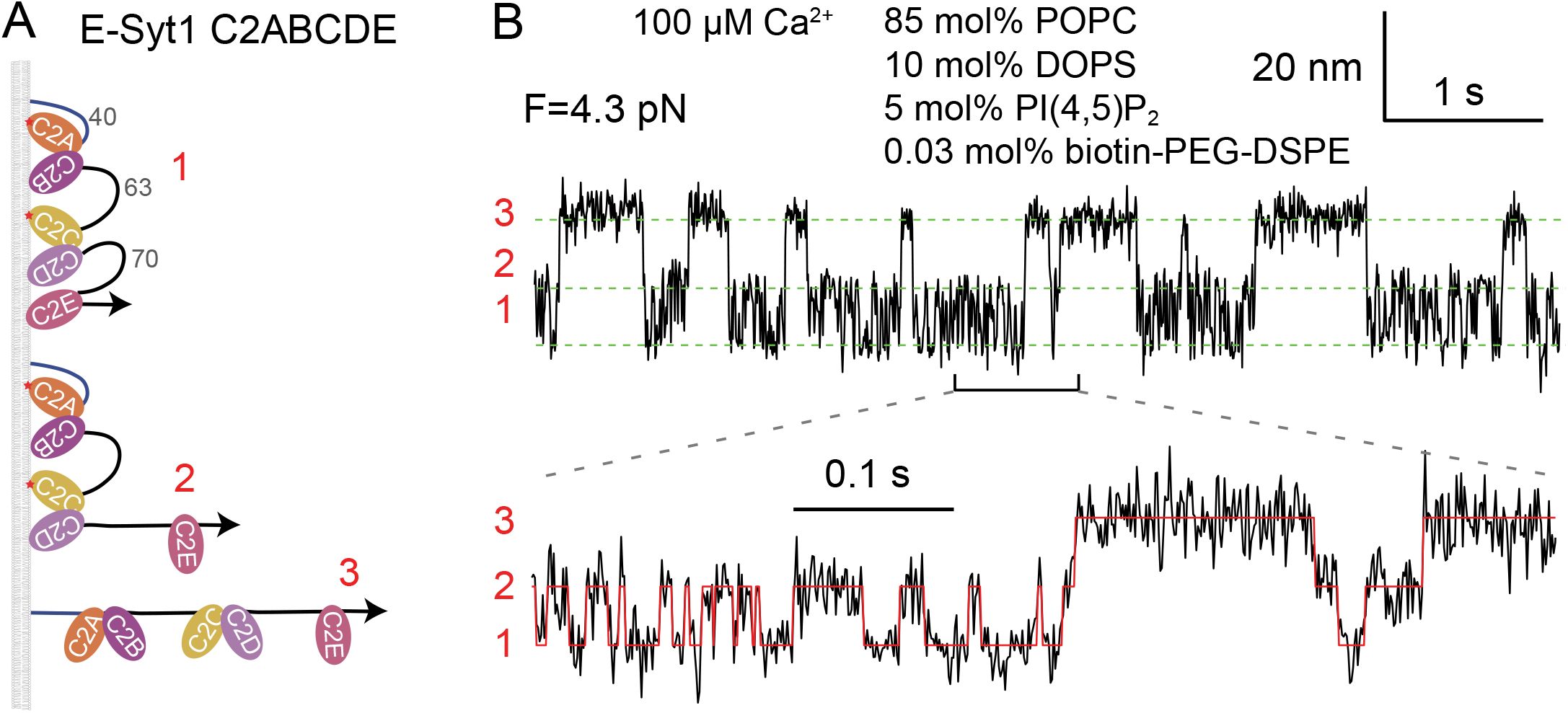
C2 domains sequentially bind to and unbind from the membrane detected by optical tweezers. A: Three C2 binding states with distinct extensions. The corresponding state numbers are shown in red. B: Time-dependent extension measured at the indicated constant mean force showing transitions of the C2 array among the different binding states. The green dashed lines mark the average positions of different states. The red curve indicates the idealized state transitions.

We develop an analytical theory to predict the *trans*-membrane binding of E-Syts required for MCS formation based on the *cis*-membrane binding parameters measured by us (Ge *et al*., 2021). The membrane binding of different C2 modules depends on the lengths of the flexible polypeptide linkers between these C2 modules. We use the Gaussian chain model to describe the conformation of each linker and extracted the membrane-binding energy and rate corresponding to each isolated C2 module. This leads to binding energy of 10.4 (±0.9) kBT for C2CD, 6 (±1) kBT for C2E in E-Syt1, 7.1 (±0.4) for C2AB, and 13.0 (±0.7) for C2C in E-Syt2 under the experimental conditions indicated in Figure 2B. Next, we calculate the *trans*-membrane binding probability of each C2 module as a function of the distance between the ER- and PM-like membranes. The *trans*-membrane binding is a tug of war between the affinity of each C2 module for the PM-like membrane and the opposing entropic force that stretches the linker between the C2 module and the ER anchor (Figure 1A). Here, the force and energy needed to stretch the linker are determined using a worm-like chain model for the linker (Marko and Siggia, 1995; Rebane *et al*., 2016). The repulsive interactions between the ER and PM membranes provide another opposing force for the *trans*-membrane binding. With an empirical potential for the membrane interactions (Zorman *et al*., 2014), the total energy of the system can be calculated to predict the equilibrium membrane distance and the probability of MCS formation.

Several salient features of the E-Syt-mediated MCSs are predicted (Ge *et al*., 2021). In the absence of Ca^2+^, E-Syt1 has less energy in the unbound state than in any C2-bound state, indicating that E-Syt1 alone cannot mediate an MCS. In contrast, E-Syt2 has minimal energy in a C2C-bound state at an equilibrium distance of ~19 nm due to the high affinity of C2C for the PM-like membrane. The free energy of this state is −2.2 kBT relative to the unbound state, consistent with a stable E-Syt2-mediated MCS. In the presence of Ca^2+^, E-Sty1 starts to tether the PM membrane at ~16 nm due to binding of both C2CD and C2E, while E-Syt2 shows a metastable state at ~11 nm due to binding of both C2AB and C2C. All these predictions, including the Ca^2+^-dependent equilibrium membrane separation, are consistent with the cellular imaging results obtained by electron and fluorescence microscopy (Chang *et al*., 2013; Giordano *et al*., 2013; Fernandez-Busnadiego *et al*., 2015; Idevall-Hagren *et al*., 2015).

### The SMP domain acts as a shuttle to transfer lipids

It is puzzling how the SMP domain transfers lipids between membranes. While many shuttle LTPs contain relatively small hydrophobic cavities that accommodate one lipid at a time (Delfosse *et al*., 2020), the SMP dimer in E-Syts forms a hydrophobic groove along its 9 nm length that can hold four phospholipids (Schauder *et al*., 2014). These observations suggest that the SMP dimer might serve as a bridge for lipid transfer. SMP belongs to the tubular lipid binding protein (TULIP) family of LTPs with conserved structural features, including long hydrophobic grooves (Kopec *et al*., 2010; Reinisch and De Camilli, 2016). One of the well-studied TULIP LTPs is cholesterol ester transfer protein (CETP) that transfers cholesterol esters and triglycerides between high-density lipoproteins (HDL) and low-density lipoproteins (LDL) in the blood (Lei *et al*., 2016). Compelling evidence suggests that CETP binds both HDL and LDL and acts as a bridge to transfer cholesterol ester or triglycerides between them (Zhang *et al*., 2012). The structural similarity between the SMP dimer and CETP also implies a bridge mechanism for lipid transfer by SMP. However, such an inference about SMP is not supported by experimental observations (Reinisch and De Camilli, 2016). The measured ER-PM distance at the E-Syt-mediated MCSs is ~20 nm in a rest state with a low cytosolic Ca^2+^ concentration and ~15 nm in an activated state with an elevated Ca^2+^ concentration (Fernandez-Busnadiego *et al*., 2015); both are significantly greater than the 9 nm length of the SMP dimer. In contrast, a recent study suggests that tricalbins, the E-Syt homologs in yeast, mediates MCSs with an ER-PM distance of 7-8 nm that are essential for lipid transfer (Collado *et al*., 2019), making it possible for the SMP dimers to form bridges at the MCSs. In summary, more evidence is required to resolve the lipid transfer mechanism by E-Syts.

It was technically challenging to distinguish whether an LTP transfers lipids as a shuttle or a bridge, due to the difficulty in controlling the membrane separation with nanometer resolution *in vitro*. Recently, Bian et al. combined the DNA-origami technique and the FRET-based lipid transfer assay to investigate the mechanism of lipid transfer by the SMP dimer (Bian *et al*., 2019) (Figure 3). They created rigid DNA rings as a scaffold to form two types of liposomes ~28 nm in diameter. The isolated SMP dimer was attached to one type of scaffolded liposomes via a disordered polypeptide linker with a specific length. These liposomes also contained NBD- and rhodamine-labeled lipids. When mixed, the two types of liposomes were joined by a rigid DNA pillar and held at a specific membrane distance. The potential lipid transfer mediated by the SMP dimer was detected via an increase in NBD fluorescence.

**Figure 3.**
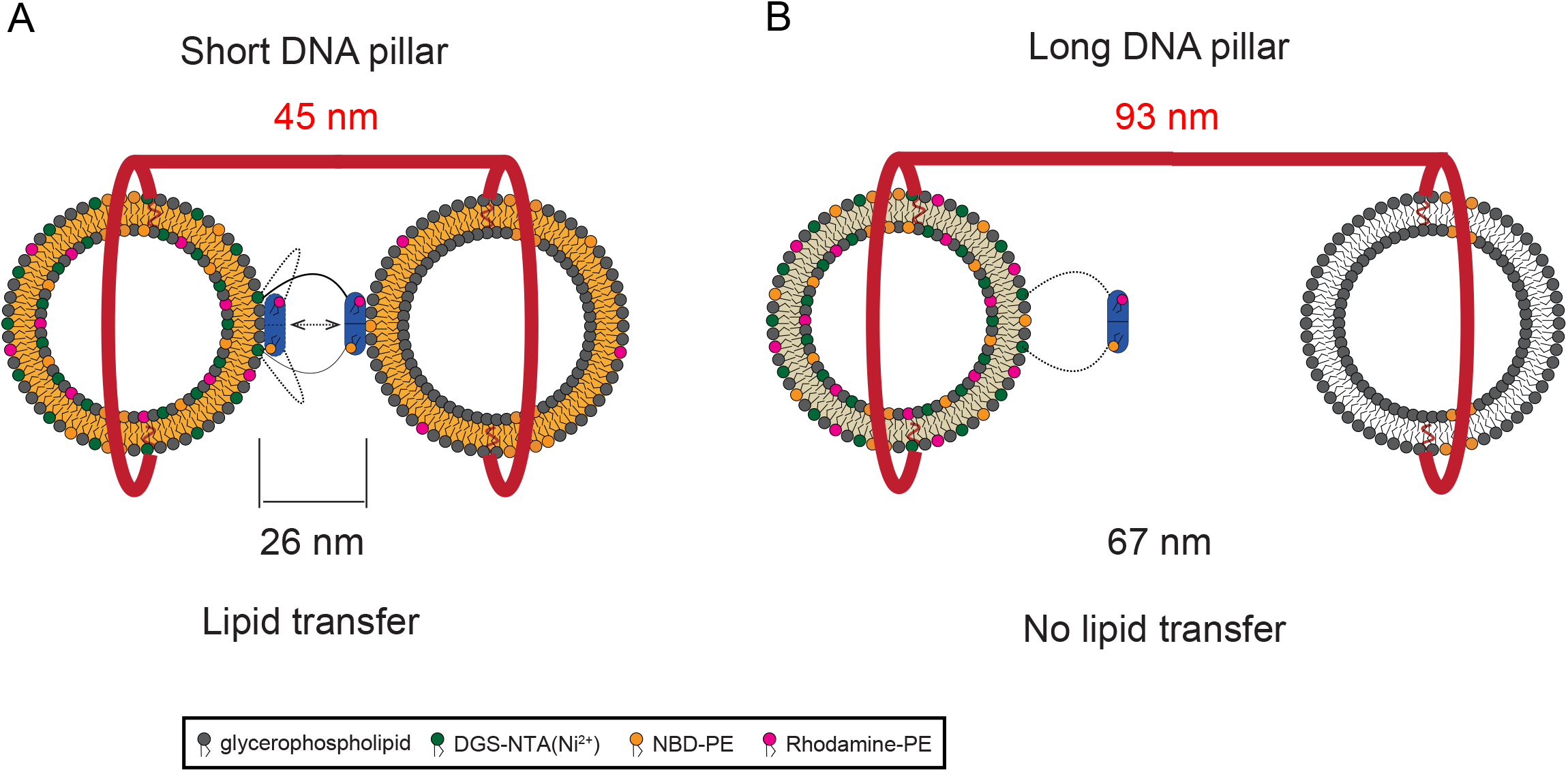
SMP dimer acts as a shuttle to transfer lipids between membranes, with their separation controlled by DNA origami. A: The SMP dimer attached to one liposomal membrane can transfer lipids to another liposomal membrane 26 nm apart that cannot be bridged by the SMP dimer (~9 nm in length). B: The short linker attaching the SMP dimer to the membrane prevents the SMP dimer from reaching another distant membrane for lipid transfer.

Using this new assay, Bian et al. demonstrated that the SMP domain of E-Syt1 transfers lipids between the two membranes even when they are separated by 26 nm (Figure 3A), significantly greater than the length of the dimeric SMP domain. Shortening the contour length of the linker from ~30 nm to ~8 nm abolishes lipid transfer, while lengthening the linker to ~85 nm rescues the lipid transfer activity, even when the two membranes are further separated to ~67 nm (Figure 3B). All these observations are consistent with a shuttle mechanism for the SMP dimer and inconsistent with the bridge mechanism under these experimental conditions. However, it remains to test whether the SMP domain can also act as a bridge to transfer lipids at a short membrane distance close to the length of the SMP dimer, given its structural resemblance to other proposed bridge LTPs.

### Lipid exchange model for shuttle LTPs

Suppose an LTP transfers lipid species 1 and 2, referred to as lipid 1 and lipid 2 for simplicity, between two membranes. Based on the lipid exchange mechanism of the Osh/ORP-family LTPs (de Saint-Jean *et al*., 2011; Mesmin *et al*., 2013; Chung *et al*., 2015; von Filseck *et al*., 2015a; von Filseck *et al*., 2015b; Ghai *et al*., 2017; Dong *et al*., 2019; Wang *et al*., 2019; Kawasaki *et al*., 2022) (Figure 4A), we proposed a reaction scheme to quantify lipid transfer (Figure 4B). We assume that the LTP is always bound by either of the lipids (see the forthcoming section: Shuttle LTPs may generally transfer lipids through the lipid exchange mechanism). When a lipid 2-bound LTP bombards lipid 1 in a membrane, the LTP exchanges for lipid 1 with a rate constant *k*_1_. Similarly, the LTP extracts lipid 2 from and inserts lipid 1 into the membrane with a rate constant *k*_2_. In the following, we will develop quantitative models for direct lipid exchange between LTPs and single membranes and LTP-mediated lipid transfer between two membranes.

**Figure 4.**
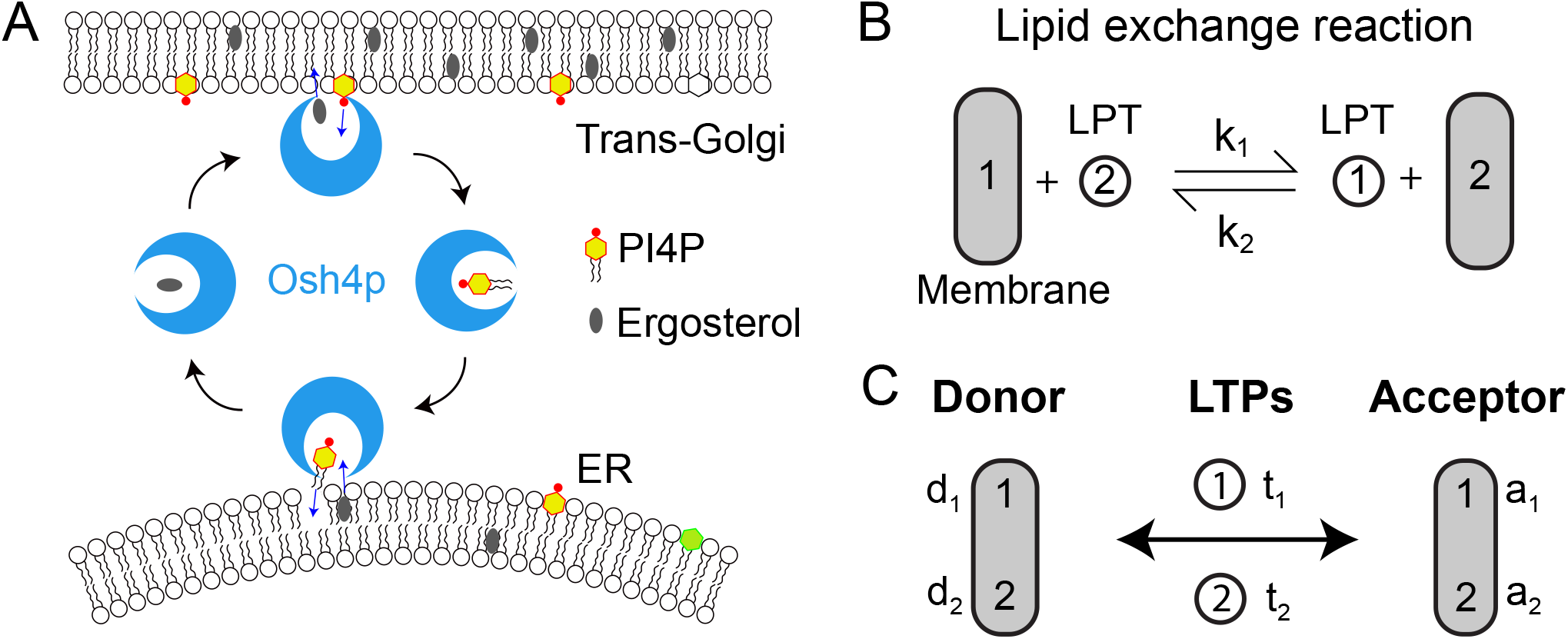
Lipid exchange scheme for lipid transport by shuttle LTPs. A: Osh4p transfers ergosterol from the ER to the trans-Golgi upgradient and PI4P downgradient in a reverse direction via a lipid exchange mechanism. B: Proposed scheme of lipid exchange between a shuttle LTP and a membrane characterized by two biomolecular reaction constants. The lipids 1 and 2 exchange between the LTP and the membrane. C: LTP-mediated lipid transfer between two membranes via two lipid exchange reactions. The concentrations of lipids 1 and 2 in the membrane and the LTP are indicated.

Direct lipid exchange between LTPs and membranes is often measured with fluorescence approaches (de Saint-Jean *et al*., 2011; von Filseck *et al*., 2015b). In these experiments, LTPs are loaded with lipid 1 and tested its exchange with an excessive of lipid 2 in a membrane by measuring the change in the fluorescence of the LTPs bound by different lipids. As a result, the effects of lipid composition and LTP mutation on the exchange rate are widely investigated. Suppose at time *t*, the membrane has molar concentrations of *d*_1_ (*t*) for lipid 1 and *d*_2_ (*t*) for lipid 2, where the number in the subscript denotes the lipid species (Figure 4C, corresponding to donor only). Correspondingly, the LTPs containing lipids 1 and 2 have molar concentrations of *t*_1_ (*t*) and *t*_2_ (*t*), respectively. Then the concentration of lipid 1-bound LTP at any time follows the law of mass action

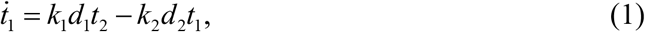

where the dot above each variable denotes its derivative with respect to time *t*. Under the experimental conditions in which the total concentrations of the two lipids in both the membrane and the LTP (*C*_1_ and *C*_2_) are much greater than the total concentration of the LTP (*C_t_*), i.e., *d*_1_ ≈ *C*_1_, *d*_2_ ≈ *C*_2_, Eq. (1) can be solved as

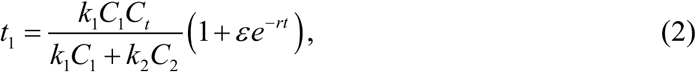

where the lipid exchange rate constant

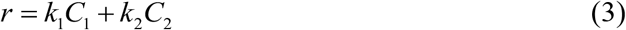

and the constant *ε* is determined by the initial concentration of lipid 1-bound LTP *t*_1_(0), i.e.,

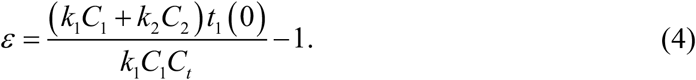

Eqs. (2)–(4) can be used to determine the exchange rate constants *k*_1_ and *k*_2_ by omitting either lipid in the membrane (*C*_1_ ≈ 0 or *C*_2_ ≈ 0) in the lipid exchange experiments. Eq. (2) indicates that the equilibrium distribution of the lipids in the LTP *t*_1_(+∞) or *t*_2_(+∞) is independent of their initial lipid distribution *t*_1_(0) or *t*_2_(0).

In the presence of another membrane, the LTP mediates lipid exchange with both membranes (Figure 4C). For the convenience of our forthcoming description, one membrane is called a lipid donor membrane and the other acceptor membrane. The two exchange rate constants may depend on membranes, as the LTP and lipids may differentially interact with proteins or non-exchangeable lipids in different membranes. Consequently, the two sets of rate constants associated with the donor and acceptor membranes may be similar or different. In the following, we will consider the two cases separately.

Our model of lipid exchange differs from the previous model developed by Drin and coworkers (von Filseck *et al*., 2015b; Ikhlef *et al*., 2021), in which several intermediate states of Osh4p-mediated lipid transfer are included, for example, the states in which Osh4p is bound to the membrane but not loaded with any lipid. Consequently, the model contains six coupled reactions with more than 10 rate constants, which make it difficult to derive an analytical solution for the lipid transfer rate and fit the model to experimental data. In contrast, our model only contains one reaction with two rate constants for similar membranes or two reactions with four rate constants for different membranes. Consequently, our model yields analytical solutions and convenient fitting to experimental data, as shown in the forthcoming sections.

### Lipid exchange between similar membranes

Suppose at time *t*, a donor membrane has molar concentrations of *d*_1_(*t*) for lipid 1 and *d*_2_(*t*) for lipid 2, while an acceptor membrane has molar concentrations of *a*_1_(*t*) and *a*_2_(*t*) for the two lipids (Figure 4C). Then the lipid and LTP concentrations at any time follow the law of mass action

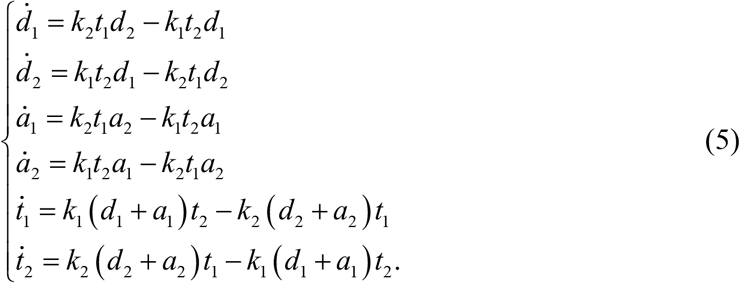

Note that these concentration variables are not independent. Based on the lipid exchange mechanism (Figure 4B), the total concentrations of the two lipids in the donor membrane (*C_d_*), the acceptor membrane (*C_a_*), and LTPs (*C_t_*), as well as the total concentration of each lipid species in both membranes and LTPs (*C*_1_ and *C*_2_), are determined by the initial experimental conditions and do not change with time, i.e.,

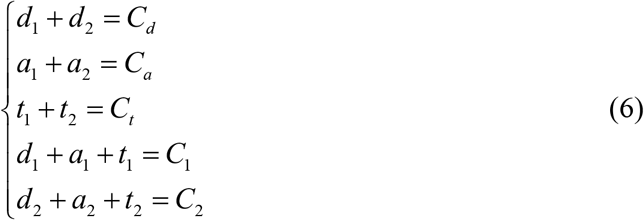

with

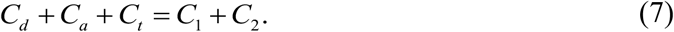

Thus, Eqs. (6) give four independent constraints and Eqs. (5) only have two independent variables. Let’s choose the fractions of lipid 1 in the donor and acceptor membranes as two independent variables, i.e.,

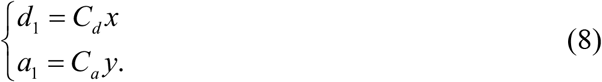

Then we can rewrite the four independent equations in Eqs. (6) as

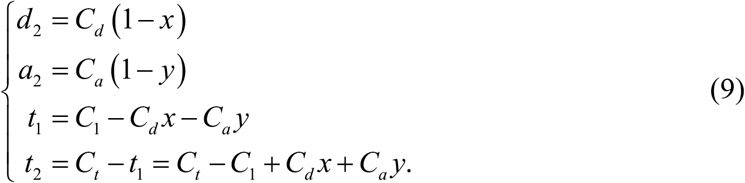

Substituting Eqs. (9) into the first two equations in Eqs. (5) yields

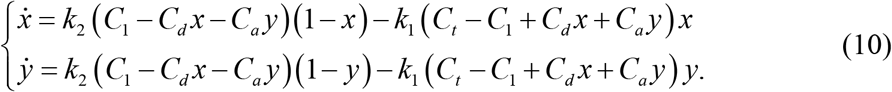

This is a system of nonlinear differential equations without exact analytical solutions available. Thus, we first numerically solved the equations under previous experimental conditions (von Filseck *et al*., 2015b). In this experiment, dehydroergosterol (DHE) and PI4P, designated here as lipid 1 and lipid 2, respectively, were transferred by Osh4p between donor and acceptor liposomes, with *d*_1_(0)=10 μM, *d*_2_(0) =0, *a*_1_(0) =10 μM, *a*_2_(0) =4 μM, *C_t_* =0.2 μM, *C*_1_ ≈ 20 μM, and *C*_2_ ≈ 4 μM (Figures 4A & 5A) or the corresponding lipid percentages listed in the middle panel of Figure 5B. The two exchange rate constants are now arbitrarily chosen to roughly match the measured lipid transfer rate (Figure 5B, top panel). The calculated DHE fractions in donor and acceptor membranes monotonically decrease and increase, respectively, and eventually equilibrate at 20/24=0.833 in both membranes (Figure 5C). In terms of Eqs. (9), the fractions of PI4P in the donor and acceptor membranes monotonically evolve to an equilibrium value of 4/24=0.167. Thus, the Osh4p-mediated lipid exchange appears to cause DHE and PI4P to mix in the two membranes.

**Figure 5.**
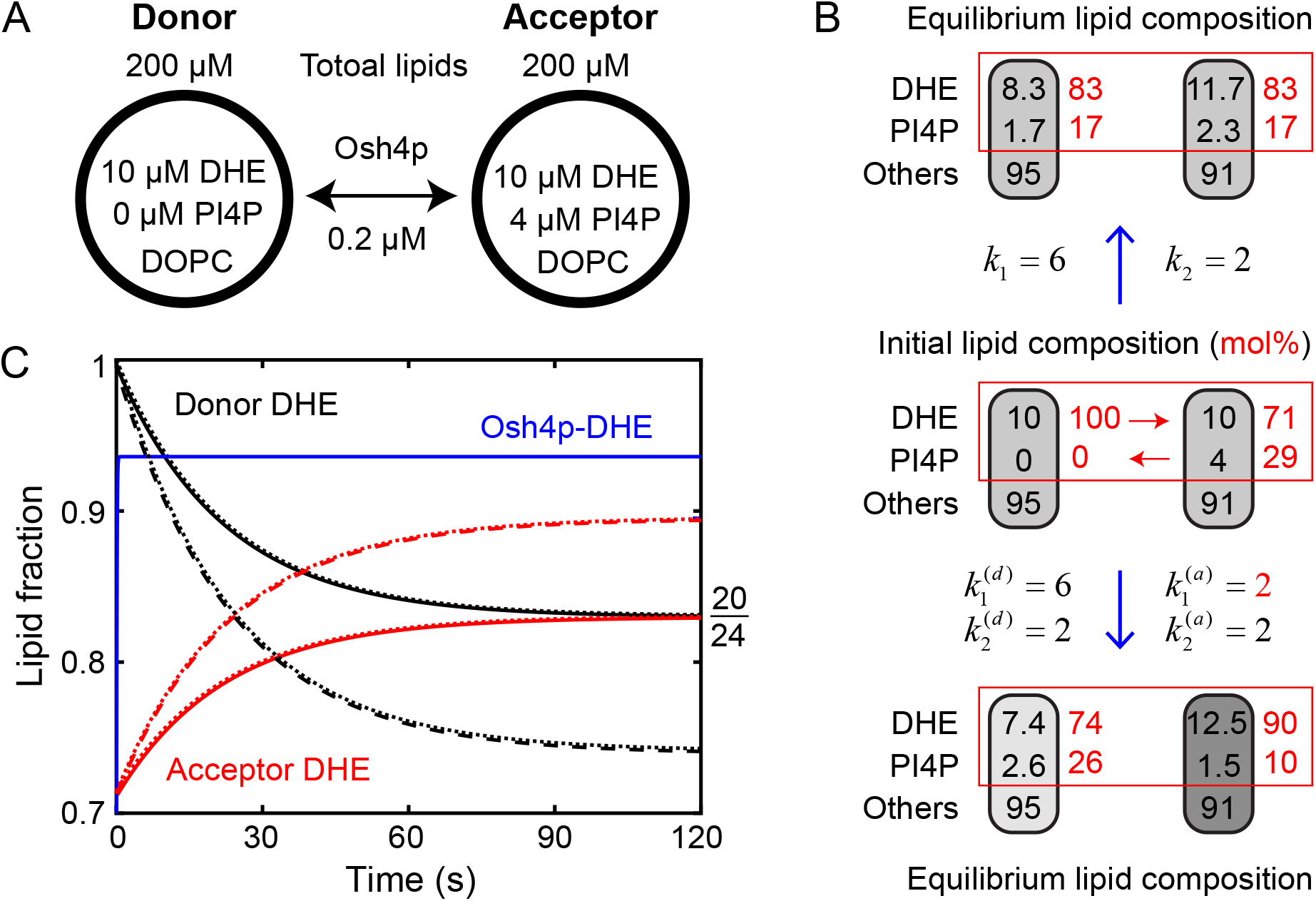
Simulations of the lipid transfer kinetics and equilibrium distributions. A: Initial concentrations of DHE and PI4P in two types of liposomes in the representative Osh4p-mediated lipid transfer assay (von Filseck *et al*., 2015b). B: Initial (middle panel) and equilibrium (top and bottom panels) concentrations of lipids accessible to Osh4p (black numbers in membranes in the unit of μM) and molar percentages within the transferable lipids (red numbers). The equilibrium lipid compositions depend on the rate constants of the lipid exchange at the two membranes (in a unit of 10^5^ M^-1^s^-1^), with equal and different membranes indicated. A gradient of the molar percentage of the exchangeable lipid drives lipid mixing, as highlighted by the red box. C: Calculated DHE fractions in the donor and acceptor membranes and Osh4p with equal (solid lines) or unequal (dashed lines) exchange rate constants (see B). The approximate analytic solutions (dotted lines) calculated based on Eqs. (12) and (22) overlap the corresponding numerical solutions. The equilibrium lipid concentrations and molar percentages within the exchangeable lipids are also shown in B.

To better understand the kinetics of lipid transfer, we derived an approximate analytic solution to the system of equations in Eqs. (10). We noticed that the fraction of DHE-bound Osh4p (Osh4p-DHE) reaches a constant within ~0.2 seconds, much earlier than the moment when the fractions of DHE and PI4P significantly change (Figure 5C). Indeed, the two lipids equilibrate among Osh4p and both membranes with a time constant of 78 ms, as calculated using Eq. (3). This observation prompted us to adopt a steady-state approximation by setting the LTP concentrations to their corresponding equilibrium values throughout the time course, i.e., 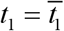 and 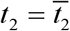, where 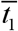 and 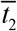 are the equilibrium concentrations of Osh4p-bound DHE and PI4P, respectively. To simplify our derivations, we further assumed that the Osh4p concentration is much less than the concentrations of the total lipids in the two membranes, i.e., *C_t_* ≪ *C_d_*, *C_a_*, as in many lipid transfer experiments (von Filseck *et al*., 2015b). Substituting these equilibrium values into Eqs. (10) and setting the right sides of the last two equations in Eqs. (5) to zero, we obtained

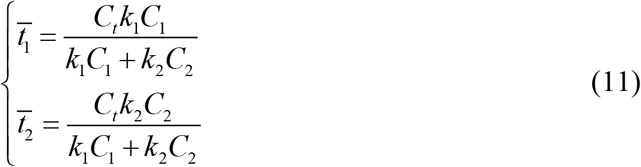

consistent with the equilibrium values calculated from Eq. (2). Here we adopted the approximations *d*_1_ + *a*_1_ ≈ *C*_1_ and *d*_2_ + *a*_2_ ≈ *C*_2_, based on the negligible LTP concentrations in the last two equations in Eqs. (6). With these approximations, we derived the analytical solutions to Eqs. (10) as

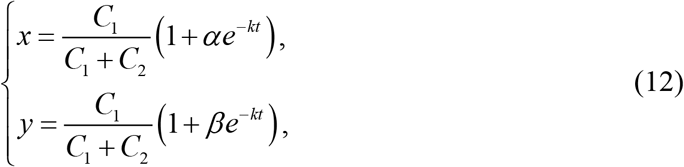

with

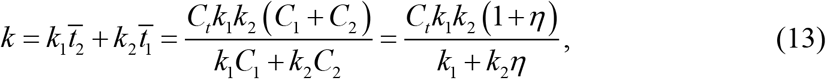

and

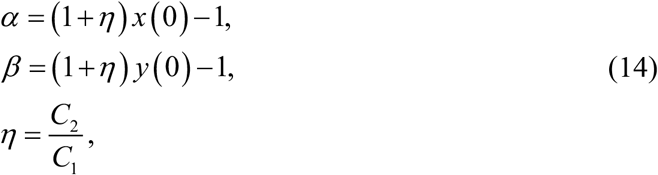

where *x*(0) and *y*(0) are the initial lipid fractions of DHE in the donor membrane and the acceptor membrane, respectively. An initial lipid transfer rate in terms of the number of lipids transferred per LTP per unit time is often measured from lipid transfer experiments (von Filseck *et al*., 2015b; Horenkamp *et al*., 2018). From Eq. (12) we calculated the initial transfer rate as

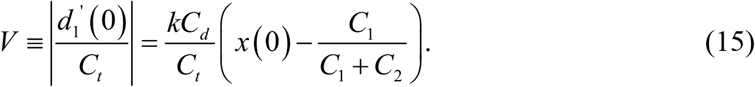

This equation shows that the initial lipid transfer rate depends on the experimental conditions as indicated by the parameters *x*(0), *C*_1_, and *C*_2_, and thus does not represent an intrinsic property of an LTP. Instead, the lipid exchange rate constants *k*_1_ and *k*_2_ better characterize the intrinsic properties of an LTP.

The analytical solution leads to interesting observations or predictions. First, despite the steady-state approximation, the analytical solutions are indistinguishable from the numerical solutions with overlapping time courses (Figure 5C), suggesting a high accuracy of the analytical solution. Second, the lipid transfer follows single-exponential kinetics characterized by a single rate constant *k* as shown in Eq. (13). This rate constant is proportional to the LTP concentration *C*_1_ and depends on both exchange rate constants *k*_1_ and *k*_2_. The former property is generally assumed in the interpretation of lipid transfer data but remains to be tested. Finally, the transfer rate also depends on the ratio of the total concentrations of the two lipids (*η*). The ratio controls the relative fraction of the LTPs bound by the two lipids and thereby the overall lipid transfer rate, as indicated by Eq. (11) and Eq. (13), respectively. The concentration ratio is a key control parameter in lipid transfer experiments *in vitro* (von Filseck *et al*., 2015b), which can be used to determine the exchange rate constants. It can be shown that the transfer rate *k* monotonically changes with *η* between the following two extreme values

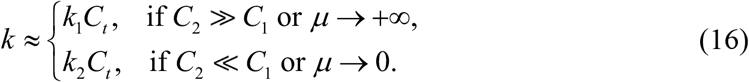

Thus, in the limiting total concentration of lipid 1 or lipid 2, the overall lipid transfer rate is limited by the extraction rate of the corresponding lipid. This property can be used to facilitate the measurement of the lipid exchange rate constants because, in these extreme conditions, the individual rate constants can be directly derived from the measured lipid transfer rate. Moreover, the transfer rate does not depend on the initial distributions of both lipids in the two membranes, such as *d*_1_(0) and *a*_1_(0), although these distributions affect the amounts of lipids transferred as shown by Eq. (14).

The analytical formula provides new insights into the mechanism of the transfer of lipids against their gradients. The solutions in Eqs. (12) give the identical fractions of the two lipids in both the donor membrane (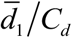 and 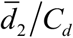) and the acceptor membrane (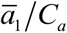 and 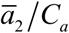), i.e.,

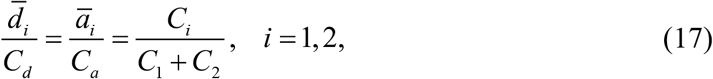

where 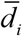 and 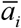, are the equilibrium concentrations of lipid 1 (*i* = 1) or lipid 2 (*i* = 2) in the donor membrane and the receptor membrane, respectively. The uniform distributions confirm that the lipid exchange causes lipid mixing. The lipid mixing is specific to the two transferable lipids, and the concentrations of nontransferable lipids do not affect the equilibrium lipid distributions within the transferable lipids, although they can change the lipid percentage within all lipids in each membrane (Figure 5). This selective lipid mixing leads to the apparent transfer of certain lipids against their gradients. For example, in the Osh4-mediated lipid transfer, the initial percentage of DHE is the same 10/105 = 9.52 mol% in both the donor and acceptor membranes. However, its percentage in the transferable lipids is 100 mol% in the donor membrane and 10/14=71 mol% in the acceptor membrane (Figure 5B, middle panel). Correspondingly, the initial percentage of PI4P in the transferable lipids is zero in the donor membrane and 4/14=29 mol% in the acceptor membrane. These different DHE and PI4P concentrations cause net transfer of DHE from the donor membrane to the acceptor membrane and PI4P in a reverse direction. Thus, both lipid transfer processes are driven by their own lipid gradients. The lipid mixing eventually leads to the same 20/24≈83 mol% for DHE and 4/24≈17 mol% for PI4P in transferable lipids in both membranes as predicted by Eq. (17) (Figure 5B, top panel). The lipid mixing dissipates any lipid concentration gradient within transferrable lipids, which differs from the antiporter ion transport mechanism, where ion gradients persist even after ion transport reaches a steady state. However, the final percentages of DHE in all lipids becomes 7.9 mol% in the donor membrane and 11.1 mol% in the acceptor membrane, leading to apparent DHE transport against its gradient. Therefore, LTP-mediated selective mixing of transferable lipids can lead to apparent transfer of lipids against their gradients.

### Lipid exchange between different membranes

The LTP-mediated lipid exchange depends on the curvature and composition of membranes (von Filseck *et al*., 2015b; Ikhlef *et al*., 2021). For example, Osh4p-PI4P extracts DHE more rapidly in DOPC liposomes than in POPC liposomes. In addition, small liposomes promote DHE extraction. To simulate lipid transfer between different membranes, we use two sets of rate constants: 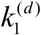 and 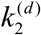 associated with the donor membrane and 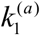 and 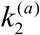 associated with the acceptor membrane (Figure 5B). Then Eqs. (5) and (10) are revised as

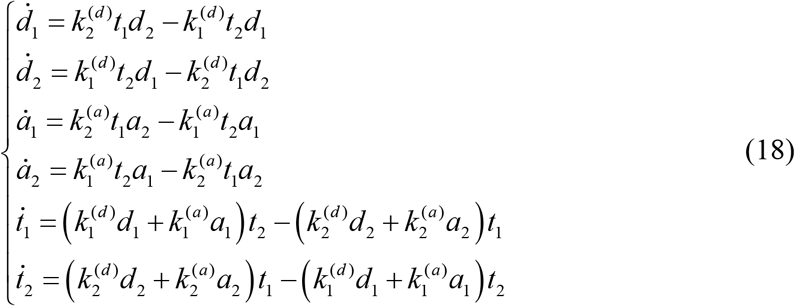

and

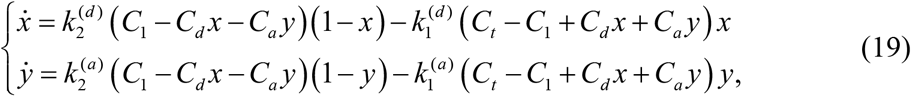

respectively. We again first numerically solved the system equations in Eq. (19) (Figure 5C). Interestingly, when we reduced the rate constant for DHE extraction from the acceptor membrane from *k* =6×10^5^ M^-1^s^-1^ to 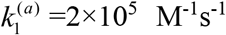, with the other rate constants unchanged (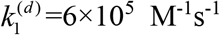 and 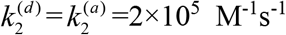), the equilibrium fractions of DHE in the two membranes become different, 70 mol% in the donor membrane and 90 mol% in the acceptor membrane (Figure 5B, bottom panel). Correspondingly, the equilibrium fractions of PI4P in the two membranes are also unequal. In this case, the free energy of the two transferrable lipids in the donor membrane (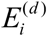, *i* = 1,2) and the acceptor membrane 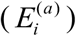 under a standard condition are different, with the energy difference

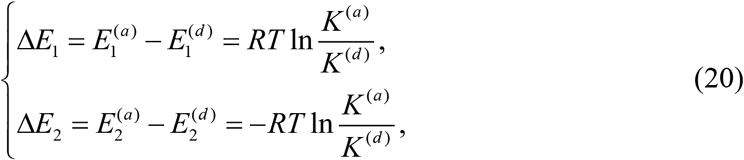

where

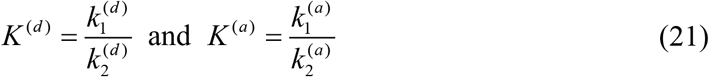

are the equilibrium constants of the lipid exchange reactions at the donor membrane and the acceptor membrane, respectively. For the different membranes simulated above, DHE has lower energy in the acceptor membrane than in the donor membrane, leading to a higher equilibrium fraction of DHE in the acceptor membrane than in the donor membrane. In this case, the LTP-mediated lipid distribution among membranes is driven by both entropy-dominated lipid mixing and energy-dominated partition, as shown by Eq. (20).

We again derived an approximate analytic solution to Eqs. (18) or (19), i.e.,

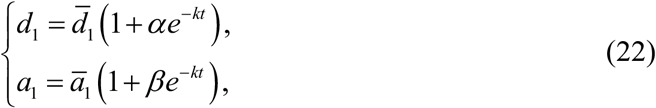

where

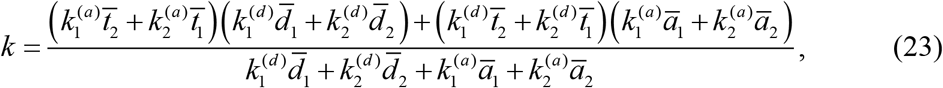

and

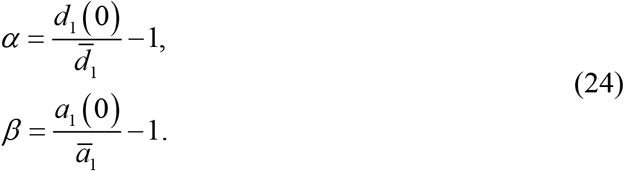

Here the equilibrium LTP concentrations are calculated as follows:

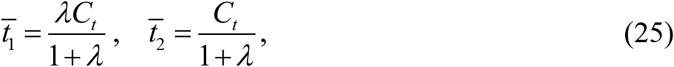

where

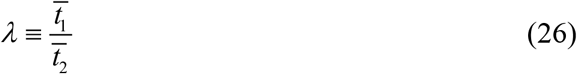

is the positive solution to the following quadratic equation

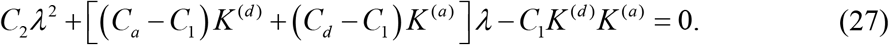

Other equilibrium concentrations are expressed in terms of 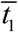 and 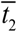 as

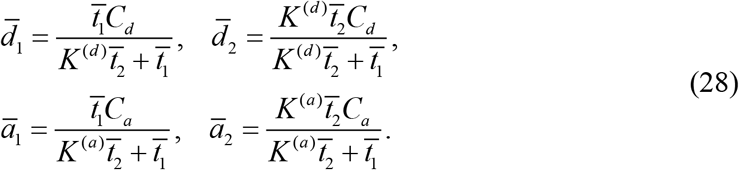

When the equilibrium constants of the lipid exchange reaction at both donor and acceptor membranes are equal, i.e., *K*^(*d*)^ = *K*^(*a*)^, the free energy of the lipids in both membranes also become equal, as is indicated by Eq. (20). In this case, all the equilibrium values shown in Eqs. (25) and (28) become equal to the corresponding values shown in Eqs. (11) and (17) under the equal membrane condition. However, the rate constant of lipid transfer expressed by Eq. (23) generally differs from the corresponding expression in Eq. (13), unless the two sets of exchange rate constants, instead of their ratios, are equal. It can be viewed as a weighted average of the lipid transfer rates calculated using the exchange rate constants of the donor and acceptor membranes (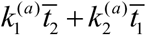 and 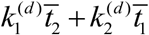) as indicated by Eq. (13).

In conclusion, we developed a quantitative model for LTP-mediated lipid transfer. We found two nonexclusive mechanisms by which LTPs transfer lipids against a lipid gradient – selective lipid mixing between similar membranes and lipid partition between membranes with different energy. Both mechanisms not only are consistent with experimental results (von Filseck *et al*., 2015b) but also lead to several interesting predictions that may be tested in future experiments to better understand the lipid transfer process.

### Lipid exchange model fit to the experimental data

To test whether the lipid exchange scheme shown in Figure 4B is consistent with the experimental results, we analyzed the time courses of Osh4p-medicated DHE and PI4P exchange measured by Drin and coworkers (von Filseck *et al*., 2015b). As shown in Figure 5A, the lipid transfer experiments began with 10 μM DHE in the donor liposomes and 0-5 μM transferable PI4P but no DHE in the outer leaflets of acceptor liposomes with the addition of 0.2 μM Osh4p. Because DOPC constitutes the majority of the total 200 μM lipids in both types of liposomes, the donor and acceptor membranes are approximately equal, and the lipid exchange reaction is characterized by two rate constants to be determined. We first calculated the time- and PI4P-dependent DHE amount transferred by Osh4p and fit the time courses with both single- and double-exponential functions (Figure 6A). We found that the transfer kinetics is not single-exponential (Figure 6A, green dashed line), but a double-exponential function fits each time course well. The double-exponential fitting revealed a fast phase with a rate (*k_f_*) of 1.2-2.8 min^-1^ (Figure 6B, bottom panel) and a slow phase with a rate (*k_s_*) of 0.15-0.20 min^-1^. The amplitude of the fast phase (*A_f_*) increases with the initial PIP4 concentration (Figure 6B, top panel), whereas the amplitude of the slow phase (*A_s_*) decreases from 1.5 μM to 0.5 μM. The fitting parameters of the fast phase lead to an initial DHE transfer rate (*V*_exp_ = *k_f_A_f_*/*C_t_*, see the definition in Eq. (15)) ranging from 7 to 22 lipids per Osh4p per minute, consistent with previous results (von Filseck *et al*., 2015b). We reasoned that the fast phase corresponds to the net transfer of DHE from the outer leaflet of the donor membrane to the acceptor membrane, while the slow phase is coupled to the flip-flop of DHE between the inner leaflet and the outer leaflet of the liposomes (John *et al*., 2002). Consistent with this interpretation, the rate constant of the slow phase is consistent with the DHE flip-flop rate constant (von Filseck *et al*., 2015b). For simplicity, we only focus on the fast phase in the following data analysis.

**Figure 6.**
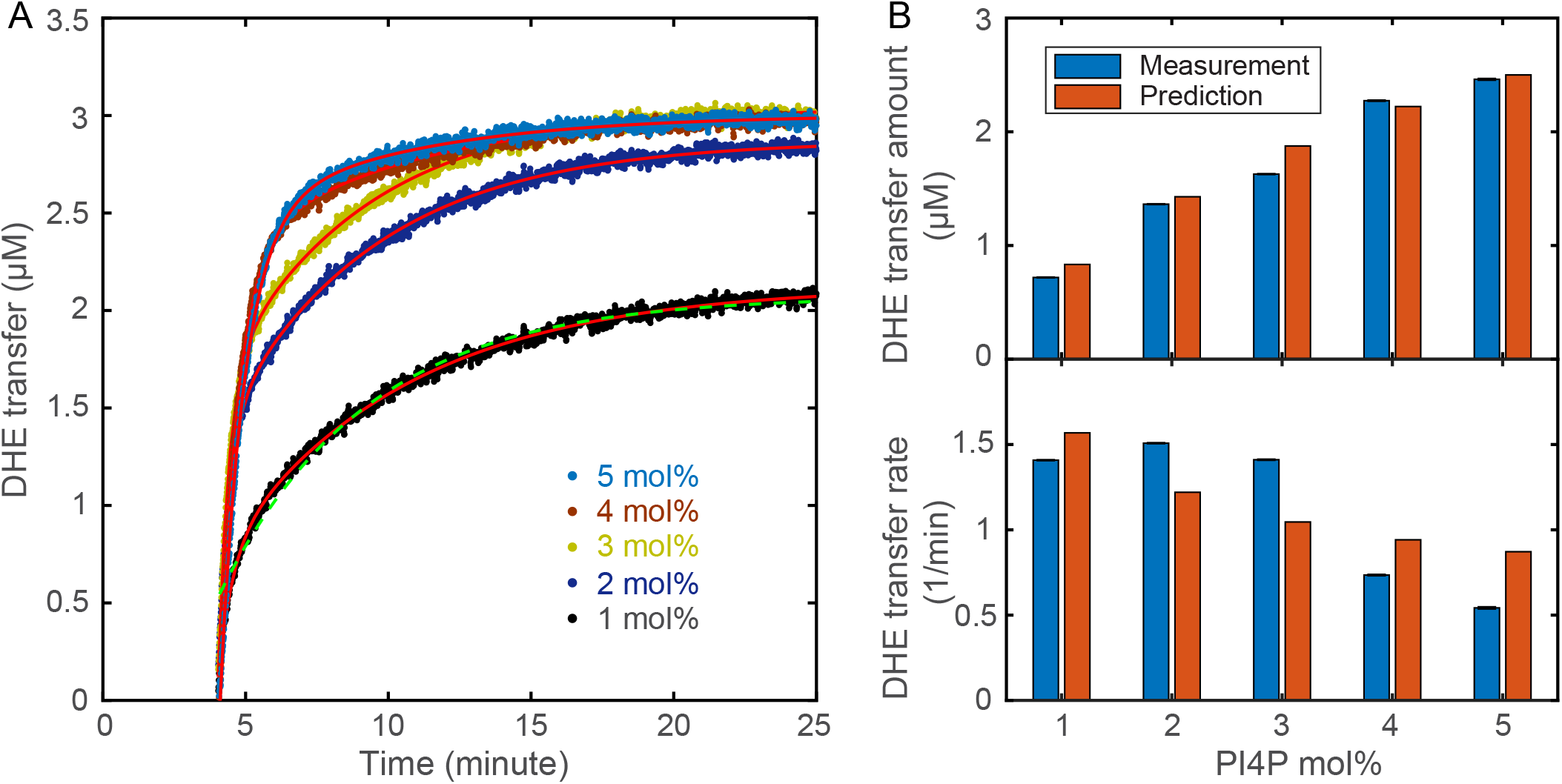
Comparisons of the measured and predicted DHE transfer amount and rates. A: Time-dependent DHE amount transferred by Osh4p (symbols) in the presence of different concentrations of PI4P in the acceptor membrane. The time courses are well fit by a double-exponential function *y*(*t*) = *A_f_* [1 −exp (−*k_f_* (*t* – *t*_0_))] + *A_s_* [1 −exp (−*k_s_*(*t* − *t*_0_))] (solid red lines) but not by a single-exponential function (green dashed line for 1 mol% PI4P). B: Comparisons of the measured and predicted DHE transfer amount (top panel) and rates (bottom panel).

The lipid exchange scheme is consistent with the measured lipid transfer kinetics. Considering the transferable lipids in the outer leaflets of both donor and acceptor membranes, we have the total lipid concentration *C_d_* = 5 μM in the donor membrane and *C_a_* = 1-5 μM in the acceptor membrane, the total DHE concentration *C*_1_ = 5 μM, and the total PI4P concentration *C*_2_ = 1-5 μM. Based on Eqs. (12) and (14) and the initial condition *x*(0) = 1, the total amount of DHE transferred by Osh4p can be calculated as

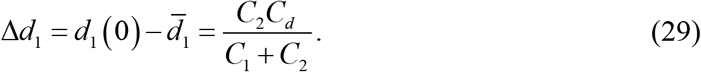

Remarkably, the predicted transferred DHE amount well matches the corresponding measurements (Figure 6B, top panel), given no fitting parameters in this case. Likewise, the initial lipid transfer rates predicted based on Eq. (15) are consistent with the measured rates. Next, we fit the calculated rates in terms of Eq. (13) to the measured DHE transfer rates at different initial PI4P concentrations. We obtained a decent fitting, which yielded the two best-fit exchange rate constants *k*_1_ = 0.2×10^5^ M^-1^s^-1^ and *k*_2_ = 4×10^5^ M^-1^s^-1^ (Figure 6B, bottom panel). Since only five data points were used to fit the two parameters, both values are regarded as estimations of the rate constants. In conclusion, the reasonable fitting of the experimental data corroborates the exchange scheme for LTP-mediated lipid transfer.

### Shuttle LTPs may generally transfer lipids through the lipid exchange mechanism

The desorption free energy needed to transfer a POPC lipid from membranes to the solution is 68 kcal/mol (27 k_B_T), with an even higher energy barrier of 96 kcal/mol (38 k_B_T) (Mclean and Phillips, 1984; Grafmuller *et al*., 2013). To efficiently solubilize the lipids required for lipid transfer, LTPs need to tightly bind lipids with unbinding energy commensurate with the desorption energy. Thus, LTPs are probably always bound by lipids, especially in the cell or at MCSs with abundant lipids. Indeed, lipids are generally found in the purified LTPs and are widely seen in their high-resolution structures (Liu *et al*., 2012; Schauder *et al*., 2014; Kumar *et al*., 2018; Valverde *et al*., 2019). Consequently, LTPs can only exchange their bound lipids with the lipids in the membrane. Since no apparent lipid transfer occurs if an LTP only recognize a single lipid species and exchanges another lipid in the same species, LTPs are expected to bind two or more species of lipids (Wong *et al*., 2017; Wong *et al*., 2019). These lipids bind to partly overlapping cavities in the LTP to accommodate both common and distinct structural features of these lipids (de Saint-Jean *et al*., 2011; Lipp *et al*., 2020). A lipid may first latch onto the distinct binding region and then displace another lipid in the overlapping binding cavity to facilitate lipid exchange. This lipid binding mode allows different lipids to exchange like toehold-mediated DNA strand displacement, which overcomes the high energy barrier for the exchange processes (Zhang and Winfree, 2009). Because the exchange process is reversible, the difference in the binding energy of both lipids for the LTP is expected to be similar to the difference in their desorption energy. Otherwise, a lipid with an excessive binding affinity for the LTP would be trapped in the binding cavity and block the exchange reaction. Consistent with this view, the LTP-lipid binding affinity appears to be tuned to a similar level for efficient exchange of the cognate lipids (Ikhlef *et al*., 2021). Therefore, based on the energetics of lipid extraction and solubilization, we hypothesize that shuttle LTPs are generally lipid exchangers. Consequently, our model for LTP-mediated lipid transfer may be broadly applicable to LTPs. More experiments are needed to test this hypothesis.

### Driving forces for directional and bulk lipid transfer through bridge LTPs

Experiments so far have revealed a slow lipid transfer via shuttle LTPs (<1 s^-1^) (von Filseck *et al*., 2015b; Wong *et al*., 2019). It is generally believed that bridge LTPs transport lipids much faster than shuttle LTPs to allow rapid membrane expansion observed *in vivo* (Kumar *et al*., 2018; Melia *et al*., 2020; Leonzino *et al*., 2021). Evidence suggests that ATG2 mediates lipid transfer from the ER to the autophagosome, a double-membrane organelle essential for macroautophagy (Maeda *et al*., 2019; Osawa *et al*., 2019; Valverde *et al*., 2019). The lipid transfer allows autophagosomes to grow from small seed vesicles into large organelles a few hundred nanometers to one micron in diameter (Sawa-Makarska *et al*., 2020; Ghanbarpour *et al*., 2021). The growth takes ~10 minutes (Axe *et al*., 2008; Xie *et al*., 2008; Schutter *et al*., 2020), requiring roughly three million lipids for a typical 400 nm autophagosome and a total lipid transfer rate of ~5,000 lipids per second (Melia *et al*., 2020). Although many LTPs could cooperate to meet the speed requirement, the limited size of the MCSs might not accommodate so many LTPs, for example, at the ER-autophagosome contact site, which necessitates a fast bridge LTP like ATG2.

Surprisingly, all bridge LTPs are found to transfer a few lipids per minute per LTP *in vitro* (Kumar *et al*., 2018; Maeda *et al*., 2019; Osawa *et al*., 2019; Valverde *et al*., 2019; Li *et al*., 2020; Hanna *et al*., 2021). The discrepancy may be caused by several limitations of current assays to detect lipid transfer by bridge LTPs, which lead to underestimations of their lipid transfer rates. First, experiments so far utilize the same FRET- and liposome-based assays to measure lipid transfer by both bridge and shuttle LTPs. These assays cannot distinguish lipid exchange and bulk lipid flow. Thus, it is possible that the lipid transfer rates measured for bridge LTPs may be mainly caused by lipid mixing rather than bulk lipid flow. Second, the bulk lipid flow requires the LTP to tether and bridge two membranes. However, the membrane bridging under *in vitro* experimental conditions may be unstable and transient, especially in the absence of other factors that facilitate the membrane tethering in the cell. Therefore, the measured lipid transfer rate is probably scaled by the limited duty cycle of the LTPs. Third, the actual number of bridge LTPs involved in membrane tethering is unknown. Consequently, the measured lipid transfer rate of a bridge LTP may no longer be proportional to the concentration of the LTP as the shuttle LTP, depending upon how the LTPs bridge the two membranes (Von Bulow and Hummer, 2020). Finally, and most importantly, current assays lack essential driving forces for bulk lipid flow present *in vivo*, as discussed below. New experimental and modeling methods are required to quantitatively understand lipid flow through bridge LTPs.

In the absence of any lipid synthesis, the bulk lipid flow through bridge LTPs causes an increase in the area of the acceptor membrane and an accompanying decrease in the area of the donor membrane. As a result, the bulk lipid flow is energetically coupled to the difference in membrane tension of the two membranes. In analogy to the roles of pressure in the water flow through a pipe and the transmembrane potential in the current through an ion channel, we propose that the membrane tension difference is a general driving force for the bulk lipid flow through a bridge LTP (Figure 7A). Membrane tension is defined as positive when the tension stretches the membrane and negative when the tension compresses the membrane. Thus, lipids flow from a membrane with lower tension to a membrane with higher tension. The lipid flow rate (*J*) is expected to be proportional to the membrane tension difference (△*σ*) or *J* = *G*Δ*σ*. The proportional constant (*G*) characterizes the conductance of the bridge in analog to ion channels. The membrane tension of the plasma membrane has widely been measured, which falls in the range of 0.001-0.1 pN/nm and varies with the cell type and growth stage (Sitarska and Diz-Munoz, 2020). The membrane tension of various organelles has not been well measured but may be in a similar range. A gradient in membrane tension can be generated during membrane biogenesis and morphology changes (Fogelson and Mogilner, 2014). For example, ER membranes form widespread networks containing membrane nanotubules attached to and pulled by molecular motors and cytoskeletons (Georgiades *et al*., 2017), causing dynamic membrane tension changes. In addition, membrane tension is intrinsically coupled to changes in membrane morphology, especially membrane curvatures. Any remodeling of membranes involving changes in their Gaussian curvatures is accompanied by membrane stretching or compression, as indicated by Gauss’s remarkable theorem (Gauss, 2005) (https://en.wikipedia.org/wiki/Theorema_Egregium). These changes in membrane tension could greatly accelerate lipid flow through bridge LTPs.

**Figure 7.**
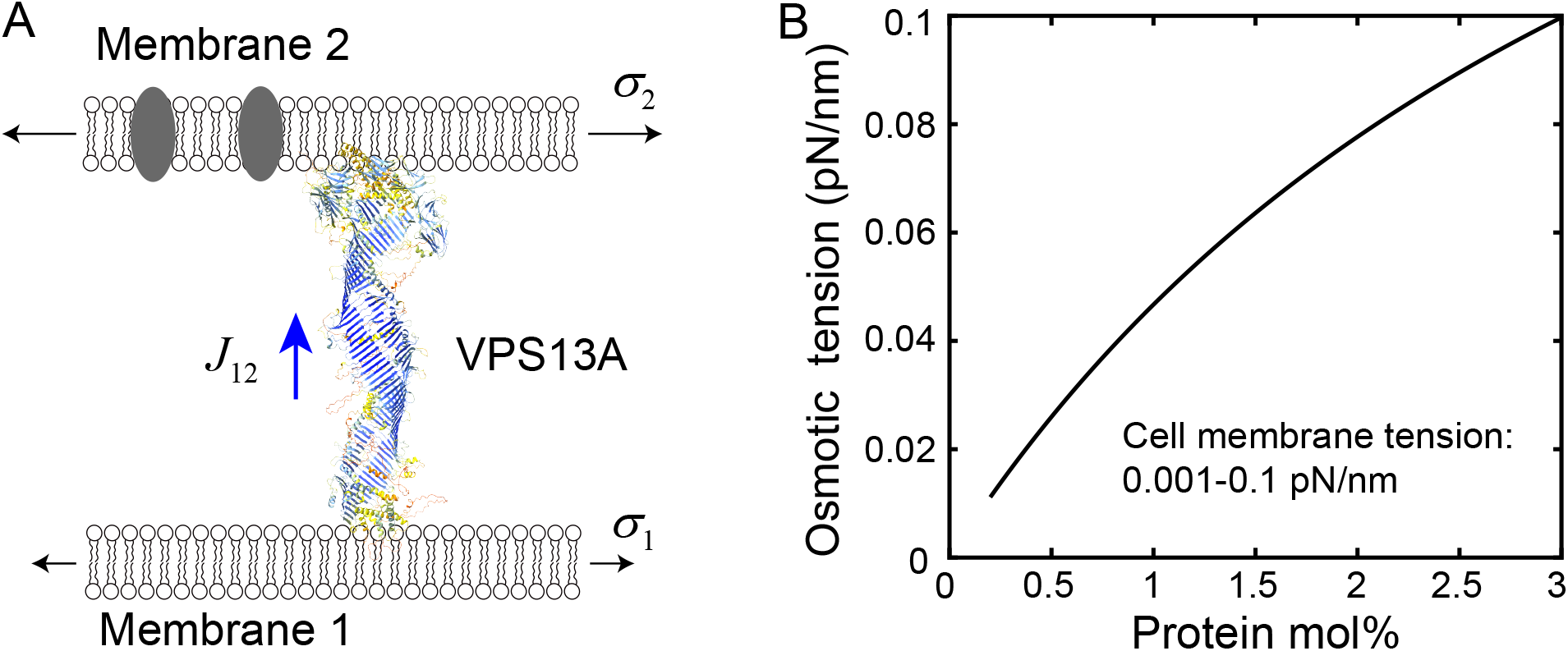
Potential driving forces for bulk lipid flow through bridge LTPs. A: Canonical membrane tension and the hypothetical osmotic membrane tension may drive lipid flow through bridge LTPs. The structure of human VPS13A, one of four isoforms of VPS13 proteins, shown here is derived from AlphaFold (Tunyasuvunakool *et al*., 2021). B: The predicted osmotic membrane tension as a function of the concentration of membrane proteins.

The lipid flow through bridge LTPs may also be driven by different lipid and protein compositions of membranes. Integral and peripheral membrane proteins are unlikely to pass through the narrow bridges (Li *et al*., 2020; Tunyasuvunakool *et al*., 2021). Consequently, an LTP bridge may act like a semipermeable membrane to generate directional lipid flow in analog to osmosis (Figure 7A). Therefore, we term this predicted new phenomenon mediated by bridge LTPs as membrane osmosis. In this case, the large molar amount of transferable lipids serves as a solvent, while the small molar amount of mobile membrane proteins and non-transferable lipids acts as a solute that cannot pass through the bridges. In parallel, we define the membrane osmotic tension (⊓) and extend the van’t Hoff formula for the three-dimensional osmotic pressure to the two-dimensional membrane tension as ⊓ = *RTc*, where *c* is the total concentration of the membrane proteins and nontransferable lipids in the membrane, with a unit of moles of total solute molecules per square meter of membrane. Membrane proteins typically constitute about half in mass or up to a few molar percent in cell membranes. Using the van’t Hoff formula, we calculated the typical membrane osmotic tension to be in the range of 0-0.1 pN/nm (Figure 7B). Interestingly, the estimated range of the osmotic membrane tension is in the same range of the membrane tension found in the cell, implying that the osmotic membrane tension may play an important role in lipid transfer and other membrane processes. Particularly, bridge LTPs may help maintain the lipid-to-protein ratio in cell membranes. Combining the two driving forces, we propose that the rate of the lipid flow through a bridge LTP from membrane 1 to membrane 2 is proportional to the difference in both canonical and osmotic membrane tension of the two membranes

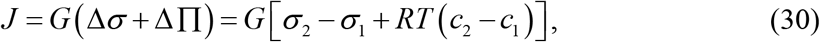

where the subscripts indicate membranes.

We estimate the lipid flow rate in the presence of a membrane tension gradient. A lipid in the LTP bridge experiences a pulling force *f* towards the acceptor membrane with higher membrane tension. This force can be estimated from the energy gain Δ*E* as the lipid passes the bridge and enters the acceptor membrane, i.e., *f* = Δ*E*/*l* = *s*Δ*σ*/*l*, where *l* is the length of the bridge and *s* = 0.7 nm^2^ the average lateral area of the lipid in the membrane (Ma *et al*., 2017). This force is balanced by a dragging force due to lipid-lipid and lipid-bridge interactions. For simplicity, we assume that the force follows the Stokes’ law, *f* = *ρv*, where *v* and *ρ* are the velocity and drag coefficient of the lipid in the bridge. We further boldly assume that the lipid has the same drag coefficient in the bridge as in the membrane, which can be estimated based on the diffusion coefficient of the lipid in the membrane (*D*) using the Einstein relation *ρ* = *k_B_T*/*D*. Combining these equations, we derive the average time taken by a lipid to pass the bridge as

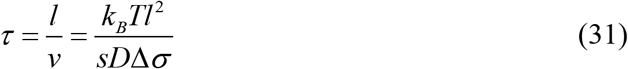

and the lipid flow rate and the bridge conductance as

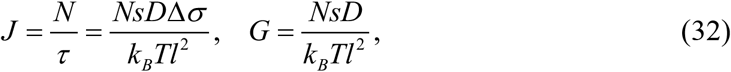

where *N* is the average number of lipids that moves together in a row in the bridge. To estimate the lipid flow properties of VPS13 and ATG2, we chose *l* = 20 nm (Valverde *et al*., 2019; Li *et al*., 2020; Tunyasuvunakool *et al*., 2021), *D* = 4 μm^2^/s (Wang *et al*., 2021b), *N* = 20 (Kumar *et al*., 2018; Valverde *et al*., 2019), and *k_B_T* = 4.1 pN×nm. In the range of the membrane tension difference Δ*σ* = 0.001 - 0.1 pN/nm, we calculated the average lipid passing time *τ* = 0.6 - 0.06 s, the lipid flow rate *J* = 34 – 3,400 s^-1^, and the conductance *G* = 3.4×10^4^ - 3.4×10^6^ nm×pN^-1^s^-1^. These calculations demonstrate that the bridge LTPs can allow fast lipid flow in the presence of physiological membrane tension.

### Implications of the driving forces in membrane biogenesis

Bridge LTPs are widely involved in biogenesis of organelles (Kumar *et al*., 2018; Valverde *et al*., 2019; Melia *et al*., 2020; Leonzino *et al*., 2021). Here we focus on the membrane expansion during autophagosome biogenesis. The dramatic membrane expansion process occurs in multiple stages and is mediated by dozens of conserved proteins (Hurley and Schulman, 2014; Gomez-Sanchez *et al*., 2021) (Figure 8A). Recent experiments suggest that the *de novo* autophagosome biogenesis starts from formation of the MCS between a small, ATG9-containing seed vesicle and the ER membrane with the help of ATG2 and ATG9 (Yamamoto *et al*., 2012; Guardia *et al*., 2020; Sawa-Makarska *et al*., 2020; Ghanbarpour *et al*., 2021). ATG9 directly binds to ATG2 and acts as a lipid scramblase to equilibrate lipids between the two leaflets of membranes at the MCS during the flow of lipids only in the outer leaflets mediated by ATG2 and other bridge LTPs (Guardia *et al*., 2020; Maeda *et al*., 2020; Matoba *et al*., 2020; Ghanbarpour *et al*., 2021). The directional lipid flow initially enlarges the seed vesicle, then likely induces its transition into a cup-shaped phagophore, a morphologically distinct precursor of the autophagosome. Further lipid transfer expands the phagophore to engulf the cytoplasmic materials to degraded, which then matures into the double-membrane autophagosome by sealing its highly curved rim. The size of phagophores is highly dynamic and adapts to the size of its cargo (Xie *et al*., 2008; Melia *et al*., 2020). Thus, the molecular mechanism of autophagosome biogenesis is key to understanding autophagy.

**Figure 8.**
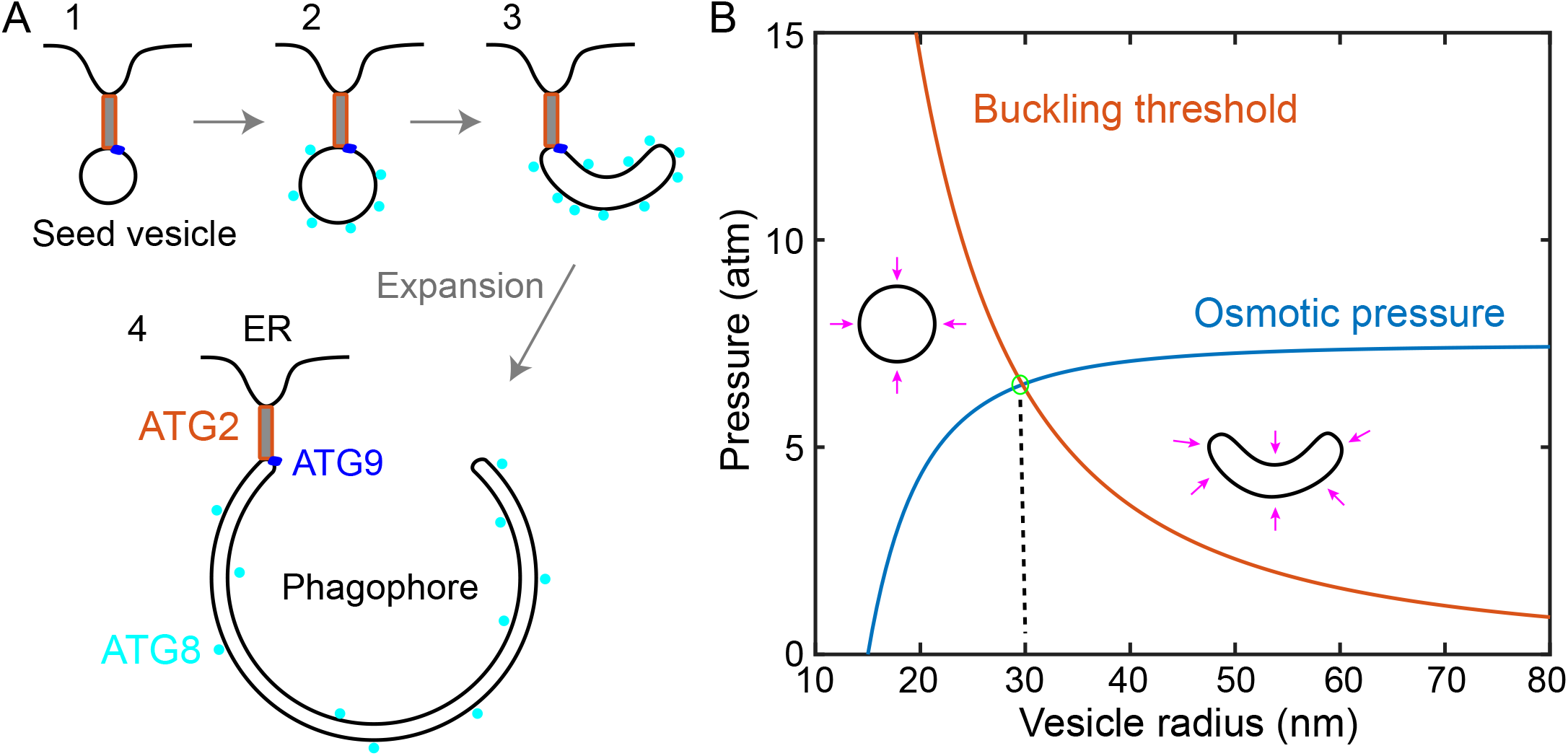
Lipid transfer, membrane conformational transitions, and osmotic pressure. A: Membrane precursors involved in autophagosome biogenesis, including the Atg9-containing vesicle tethered to the ER membrane by ATG2, ATG9, and other proteins (state 1), the growing vesicle containing lipidated ATG8 (state 2), the cup-shaped phagophore (state 3) with its convex side attached to cytoplasmic materials to be degraded (not shown), and the expanded phagophore (state 4) engulfing cytoplasmic materials. Sealing of the top rims of the expanded phagophore via membrane fission leads to a mature phagophore. B: Comparison of the vesicle buckling pressure and osmotic pressure as a function of the vesicle radius. A growing spheric vesicle buckles into a cup-shaped vesicle as the compression osmotic pressure reaches the buckling pressure at 6.5 atm and a vesicle radius of ~30 nm (marked by a green circle). The pressure unit is atmospheric pressure (atm). For the calculations, the osmolarity of cytosol is chosen as 300 mOsm. The osmolarity inside the seed vesicle 15 nm in radius is the same but decreases as its radius increases.

Various factors in the cell may generate the canonical and osmotic membrane tension to regulate autophagosome biogenesis. Cytoskeletons and their associated motors play wide roles in biogenesis and remodeling of organelles, for example, in endocytosis, and help regulate membrane tension of the PM (Shi *et al*., 2018). Likewise, cytoskeletons are essential for nearly every stage of autophagosome biogenesis (Kruppa *et al*., 2016; Kast and Dominguez, 2017). Agents that block actin or microtubule polymerization inhibit phagophore growth. Although the underlying mechanisms are generally unclear, it is possible that cytoskeletons and their associated molecular motors attach to and pull on growing phagophores, which induce tension in phagophore membranes that draw lipids from the ER for phagophore membrane expansion via ATG2.

The ATG2-mediated lipid flow may also be driven by the osmotic membrane tension induced by many peripheral proteins essential for autophagosome growth, especially ATG8 (Melia *et al*., 2020; Sawa-Makarska *et al*., 2020). ATG8 is covalently and reversibly conjugated to phosphatidylethanolamine (PE) in phagophore membranes during autophagosome biogenesis (Xie *et al*., 2008; Abdollahzadeh *et al*., 2017). The lipidated ATG8 is the most abundant protein on phagophore membranes and long serves as a defining marker of autophagosomes (Geng *et al*., 2008). Importantly, the amount of lipidated ATG8 correlates with the size of autophagosome for an unknow mechanism (Xie *et al*., 2008). We hypothesize that the concentration of lipidated ATG8 contributes most to the osmotic membrane tension of phagophores, which in turn regulates lipid flow to phagophore by ATG2, in addition to the many other functions of ATG8 lipidation in autophagy (Shpilka *et al*., 2011). In yeast, each autophagosome has an average radius of ~150 nm and contains 272 lipidated ATG8 proteins under growth conditions (Xie *et al*., 2008). It appears that these ATG8 proteins accumulate in the early phase of phagophore growth. Assuming the same number of lipidated ATG8 on a spheric vesicle with a radius of 30 nm or 150 nm, we estimated ATG8 generated osmotic membrane tension of 0.1 pN/nm or 0.002 pN/nm, respectively. Therefore, the small vesicle probably has much higher osmotic membrane tension than the ER, driving fast lipid flow from the ER to the vesicle. In contrast, the mature autophagosome bears osmotic membrane tension similar to or even lower than that of the ER, which prevents further growth of the autophagosome. Thus, the dilution of the lipidated ATG8 accompanying autophagosome growth may provide a simple mechanism to regulate the autophagosome size via the osmotic membrane tension.

Directional lipid flow is intimately coupled to membrane configurations. The lipid flow may help generate the characteristic cup-shape phagophore essential for engulfing the cytoplasmic materials to be degraded. This coupling between lipid transfer and membrane configurations is recently proposed by Ghanbarpur et al. based on conservation of the vesicle volume (Ghanbarpour *et al*., 2021), as is demonstrated in previous studies on membrane mechanics and morphology (Seifert, 1997; Flatt and Bruce, 2009; Bahrami *et al*., 2017). Here we clarify this important concept based on vesicle buckling induced by the osmotic pressure. Except for ATG9, the phagophore membrane is essentially devoid of any integral membrane proteins (Fengsrud *et al*., 2000), including ion channels and transporters that may regulate the osmotic pressure of phagophores. The cytosol of a cell typically has an osmolarity of ~300 mOsm, which could generate a maximum compression pressure of 7.5 atmospheric pressure (atm) on a membrane bound organelle corresponding to zero osmolarity in its interior. If the mechanical strength of the organelle could not withstand the pressure, the organelle would flatten, eventually turn into a cup-shape with smaller volume to minimize both the osmotic pressure and membrane bending energy (Seifert, 1997).

We predict a transition in membrane configuration from a sphere to a cup-shape during phagophore expansion. To this end, we calculated the osmotic pressure of a spherical vesicle and its buckling threshold as a function of the vesicle radius (Figure 8B). Any thin spherical shell buckles at a maximum compression pressure

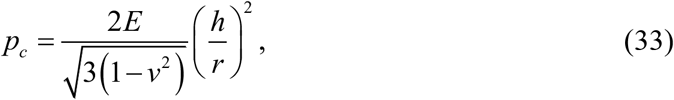

where *h* and *r* are the thickness and radius of the shell, respectively, and *E* and *v* are the Young’s modulus and Poisson’s ratio of the material of the shell (Abbasi *et al*., 2021). For lipid bilayers, we chose *h* = 5 nm, *E* = 19.3 MPa or 193 atm (Picas *et al*., 2012), and *v* = 0.25 (Jadidi *et al*., 2014). To calculate the osmotic pressure of the vesicle, we assumed that the amount of membrane impermeable solutes inside the vesicle remains constant as the vesicle grows and the initial amount is determined by the seed vesicle with zero osmotic pressure. Thus, the osmotic pressure (*P_osm_*) increases as the vesicle grows, i.e.,

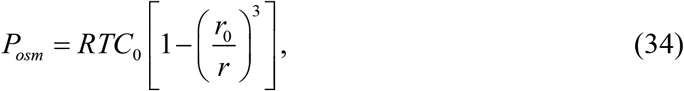

where *C*_0_ = 300 mOsm is the cytosol osmolarity, *r*_0_ = 15 nm the radius of the seed vesicle (Yamamoto *et al*., 2012; Hurley and Schulman, 2014), and *r* the radius of the growing vesicle. Eqs. (33) and (34) shows that, as the vesicle radius increases, the buckling pressure decreases while the osmotic pressure increases. At a critical radius of ~30 nm, the compression osmotic pressure reaches the buckling pressure of the vesicle and vesicle buckles into a cup shape. Once the buckling occurs, the vesicle volume decreases to reduce the osmotic pressure. Because a buckled vesicle, especially a large one, bears little compression pressure, its interior volume may eventually return to a volume close to that of the seed vesicle. Although the membrane configurational transition reduces the energy associated with osmotic pressure, the highly curved phagophore membrane increase the energy barrier for lipid transfer, thereby impeding continuing lipid flow. This may also explain why the lipid transfer rates measured for the bridge LTPs *in vitro* are much smaller than expected. In the cell, many peripheral proteins involved in autophagy, including lipidated ATG8, may bind phagophores to stabilize their curved membranes (Hurley and Schulman, 2014; Bahrami *et al*., 2017; Nguyen *et al*., 2017; Melia *et al*., 2020), facilitating the configurational transition and continuing lipid flow via bridge LTPs.

In conclusion, the gradients of canonical and osmotic membrane tension may drive the directional flow of bulk lipids via bridge LTPs required for organelle biogenesis. The lipid flow is intimately coupled to membrane configuration and osmotic pressure and is facilitated by various proteins that pump lipids in an ATP-dependent manner, remodel membranes, regulate the volumes or osmotic pressure, and equilibrate the lipids between the two leaflets of bilayers. (Owens *et al*., 2019; Matoba *et al*., 2020; Ghanbarpour *et al*., 2021).

## Conclusions and Perspectives

Despite extensive studies, there have been no tractable quantitative models for lipid transport mediated by LTPs. We have developed an analytic theory for the lipid transfer process mediated by shuttle LTPs based on a simplified lipid exchange scheme. The theory is consistent with some experimental results and accounts for the widely observed transport of lipids against their gradients via two nonexclusive mechanisms: selective lipid mixing between similar membranes and partitioning between different membranes. The theory also leads to many interesting predictions awaiting further experimental tests. First, we found that the initial lipid transfer rate often used to characterize an LTP does not represent an intrinsic property of the LTP and may not be compared among different LTPs tested under different experimental conditions. Instead, the lipid exchange rate constants better characterize the intrinsic properties of an LTP. Second, the lipid transfer rate is proportional to the LTP concentration. Finally, the lipid transfer rate depends on the relative concentrations of the two exchangeable lipids (*η*), but not their initial distributions among the two membranes. In addition, the membrane-dependent rate constants for lipid exchange should be examined. A systematic test of these predictions will not only deepen our understanding of the lipid transport mediated by the shuttle LTPs but also confirm or extend the quantitative models.

We proposed two potential driving forces for the bulk flow of lipids through bridge LTPs. Despite compelling evidence for VPS13 and ATG2 as bridge LTPs, direct support for the bridge model has been lacking. Improved assays will be required to monitor bulk lipid flow and the accompanying membrane area changes. These assays may need controlled canonical or osmotic membrane tension, membrane tethering, and osmotic pressure. In addition, new assays may be needed to better distinguish between shuttle and bridge mechanisms of an LTP. Some LTPs might transfer lipid as both shuttles and bridges, at least *in vitro*, depending upon experimental conditions. An N-terminal domain of ATG2 transfers lipids and can substitute the wild-type ATG2 in autophagy in the cell (Valverde *et al*., 2019). An ATG2 mutant that fails to tether membranes still transfers lipids, albeit with one-third of the activity of the wild-type (Maeda *et al*., 2019). These observations seem to corroborate a shuttle mechanism of lipid transfer by at least the mutant ATG2. Finally, the quantitative models for lipid transfer and membrane contact formation should be combined to simulate lipid transfer at the MCSs.

In conclusion, continuing discoveries of new LTPs and their biological functions call for quantitative models for lipid transport and membrane contact formation regarding their kinetics and thermodynamics. These models will also improve our understanding of the thermodynamics of membranes and potential new mechanisms that govern organelle biogenesis and stability.

## Acknowledgments

We thank Pietro De Camilli, Thomas Melia, and Karin Reinisch for reading the manuscript, and Guillaume Drin for providing the lipid transfer data shown in Figure 6.

